# KBoost: a new method to infer gene regulatory networks from gene expression data

**DOI:** 10.1101/2021.04.01.438059

**Authors:** Luis F. Iglesias-Martinez, Barbara De Kegel, Walter Kolch

## Abstract

Reconstructing gene regulatory networks is crucial to understand biological processes and holds potential for developing personalized treatment. Yet, it is still an open problem as state-of-the-art algorithms are often not able to process large amounts of data within reasonable time. Furthermore, many of the existing methods predict numerous false positives and have limited capabilities to integrate other sources of information, such as previously known interactions. Here we introduce KBoost, an algorithm that uses kernel PCA regression, boosting and Bayesian model averaging for fast and accurate reconstruction of gene regulatory networks. We have benchmarked KBoost against other high performing algorithms using three different datasets. The results show that our method compares favorably to other methods across datasets. We have also applied KBoost to a large cohort of close to 2000 breast cancer patients and 24000 genes in less than 2 hours on standard hardware. Our results show that molecularly defined breast cancer subtypes also feature differences in their GRNs. An implementation of KBoost in the form of an R package is available at: https://github.com/Luisiglm/KBoost.

## 1 Introduction

Gene regulatory networks (GRNs) are models that describe how transcription factors (TFs) orchestrate the expression of other genes [1-3]. In GRNs the nodes are TFs and genes, and the edges represent regulatory interactions. TFs bind to the promoters of genes to activate or silence the production of mRNA [4]. By doing so, TFs help control gene expression and thus modulate or enable cellular processes [5]. Most existing approaches of studying gene regulation focus on the effect of a single TF on the rest of the genes. However, TFs can regulate the expression of other TFs and thus a perturbation of a single TF can propagate information throughout the whole system. The potential for a single TF to control a system depends on certain network features: a TF that regulates many genes will obviously have a large influence, but the contrary is not necessarily true. Even if a TF regulates only a small number of genes directly, it can still play a major role in the system if these genes in turn regulate many others. This is the concept of closeness centrality in graph theory. Therefore, understanding a system’s GRN architecture can reveal the TFs that are important for a specific phenotype and thus can explain molecular pathogenesis and reveal potential drug targets.

Several methods have been developed to predict GRNs using gene expression data [2, 3, 6-9]. These methods formulate the problem of inferring a GRN as an unsupervised classification problem. Typically, a GRN is formulated as a weighted graph, trained on unlabeled data, whose edge values represent the predicted probability that each TF regulates other genes (including other TFs).

Several groups have used different algorithms based on different mathematical formulations to infer GRNs from gene expression data. These include Bayesian networks, correlation metrics, mutual information methods and parametric and non-parametric regression. A seminal paper published in 2012 showed that correlation, mutual information and Bayesian networks tended to perform far worse than methods based on regression [3]. For this reason, in this work we focused only on regression based GRN inference methods.

Regression based GRN inference methods build a mathematical model of the expression levels of a target gene given the expression levels of different TFs. The model goodness of fit is used to estimate the probability that the TFs in a model regulate the target gene. That is, if the expression of a TF poorly predicts the expression of a gene, then it is not likely to regulate it and *vice versa*. Some of the popular regression based GRN inference algorithms are summarized in Fig. 1.

**Figure 1.**
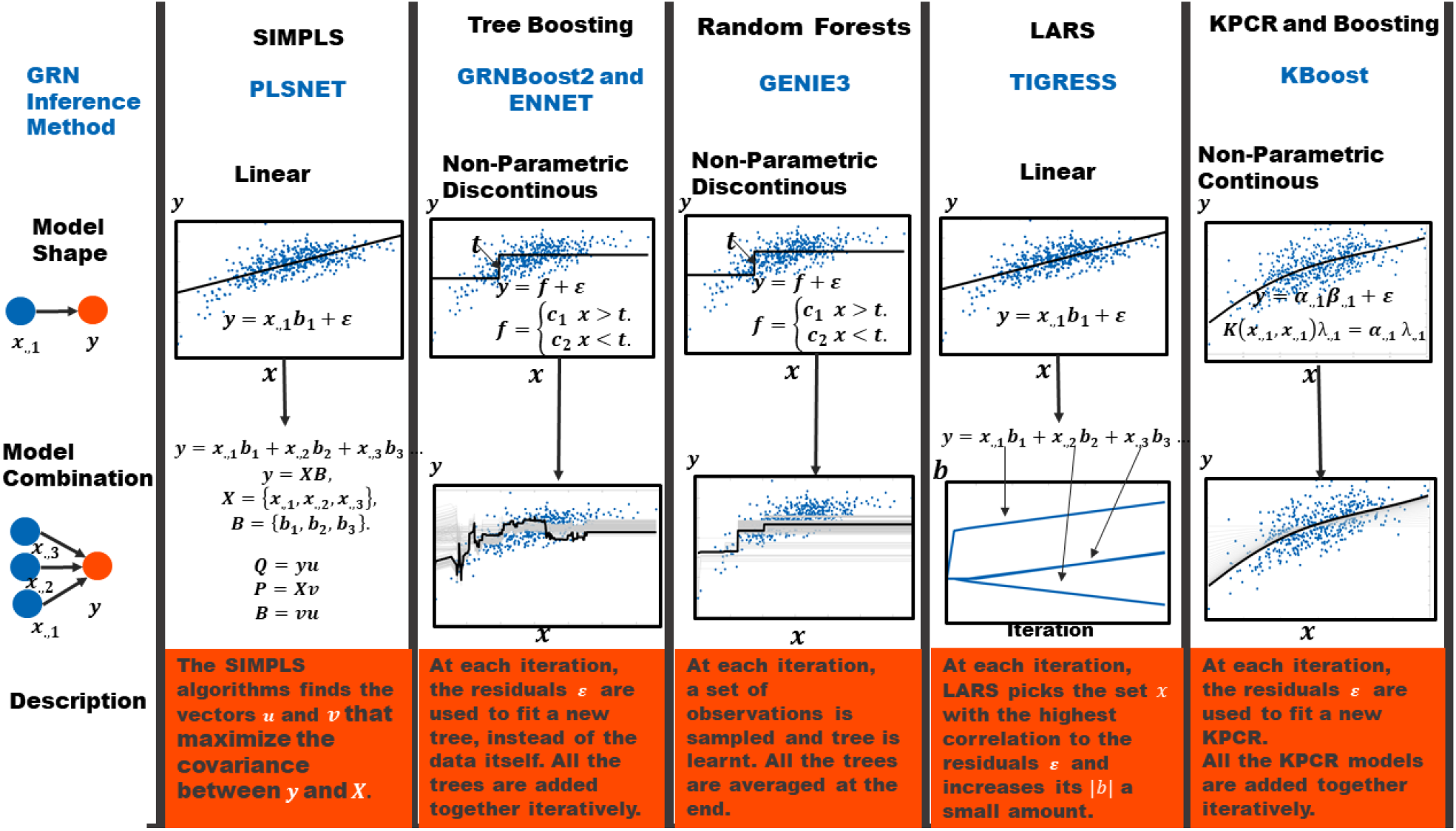
Regression-based Gene Regulatory Network Inference Algorithms. Schematic describing how different regression based GRN inference methods work. These methods are based on different machine learning algorithms. We show 6 methods based on different machine learning algorithms that differ on the model shape and the way models for different TFs are combined.They were selected because they represent major types of machine learning methods used for GRN reconstruction and because of their high performance in the DREAM 4 and DREAM 5 challenges. PLSNET uses partial least squares and fits a linear model between TFs and targets. TIGRESS uses a linear model with different lasso parameters They both rely on the assumption that the expression of a gene is proportional to the expression levels of the TFs that regulate it. GRNBoost2 and ENNET use boosting to learn different tree models between TFs and targets. GENIE3 also uses tree models, however they iteratively resample different subsets of observations and potential TFs per target and create an ensemble of tree models. Unlike linear models, tree models do not rely on any assumption between the relationship of a TF and a target, however they are not continuous models. Finally, we propose KBoost which uses boosting and KPC regression to model the relationship between TFs and targets. KPC regression can fit any relationship between TFs and targets plus it produces a continuous model.

Parametric GRN inference algorithms use a mathematical formulation that describes gene regulation using a predefined expected form. The most popular algorithms belonging to this class are linear regression based. In other words, they assume that the expression of a target gene is a linear combination of the expression levels of the TFs that regulate it [8-11]. PLSNET and TIGRESS both use a linear regression approach [8, 9]. PLSNET uses a form of partial least squares known as SIMPLS [8], while TIGRESS uses the least angle regression algorithm (LARS) [9]. LARS is an effective approach for obtaining different solutions for the linear regression coefficients with different absolute norms [12]. This algorithm takes an iterative approach: at each step, the residuals between the response variable and the linear regression model are compared with the explanatory variables, and because the models are trained iteratively on the residuals it can be described as a boosting algorithm [12]. TIGRESS couples LARS with stability selection and uses the frequency with which each TF is chosen in iterations of LARS as a proxy for the predictive probability that a TF regulates a gene [9]. The effectiveness of linear model-based approaches relies on how well a target gene’s expression levels can be approximated using linear functions.

For this reason, nonparametric regression methods have been proposed. In particular, regression trees have been very successful for GRN inference in different datasets [2, 6, 7]. Regression trees separate the explanatory variables into regions and give the response variable a value depending on what region of the explanatory variables each observation resides in [13, 14]. The GENIE3 method uses the regression version of the random forest algorithm [2, 13] - this algorithm iteratively samples different observations and explanatory variables and builds regression trees with a fixed depth. The different trees’ predictions are then combined to form the final prediction [13]. GENIE3 runs the random forest algorithm once per target gene. At each step, the variable importance of each TF is calculated as the proportion of explained variance of the target gene. The GENIE3 algorithm won two important GRN inference challenges, namely the DREAM 4 multifactorial sub-challenge and the DREAM 5 challenge [2]. The datasets for these challenges are now widely used benchmarks to test the performance of different GRN inference algorithms. Besides the random forest approach, regression tree models can also be trained using a stochastic gradient boosting approach. In the context of GRN inference, ENNET and GRNBoost2 are based on regression tree stochastic gradient boosting [6, 7]. Stochastic gradient boosting is a form of a gradient based function optimization applied on model space [14]. Very generally speaking, tree boosting is similar to LARS except that it uses tree models instead of linear models [12].

Here we present KBoost, a method that uses kernel PCA (Principal Component Analysis) regression and gradient boosting to reconstruct gene regulatory networks. Kernel PCA regression (KPCR) is a nonparametric regression technique that does not require the data to follow a strict form [15, 16]. This is accomplished by transforming the explanatory variables, in our case the TF expression levels, using a kernel function. The kernel function yields a symmetric matrix that corresponds to the dot product of nonlinear features that are a function of the explanatory variables. For many kernel functions these implicit nonlinear features correspond to polynomials of the explanatory variables [17]. In fact, the radial basis kernel function (RBF) is the dot product of an infinite number of polynomials without having to calculate them explicitly [18]. Using polynomials for regression is an attractive approach, as according to the Weierstrass approximation theorem any continuous real function can be uniformly approximated by polynomials. Furthermore, kernel PCA lets us obtain the principal components of this large number of polynomials by using the kernel matrix with much smaller dimensions. Thus, with KPCR we can use the first principal components of a large number of implicit features to approximate any continuous function - in our case the relationship between a TF and a target gene. Gradient boosting is used to iteratively combine predictions from different models, in our case KPCRs from different TFs [7, 14]. Unlike most GRN inference methods, we use a greedy model search that allows us to drastically reduce the running time. KBoost uses a Bayesian formulation, and thus we can include information from previous experimental data and increase the accuracy of our predictions [11, 19]. KBoost performed better than other algorithms on several benchmark datasets.

## 2 Methods

### 2.1 KBoost Overview

The aim of KBoost is to provide a fast and scalable algorithm for the accurate inference of GRNs (Fig. 2). It takes mRNA expression (microarray or RNA-seq) and, optionally, previously found TF-target interactions (e.g. ChIPseq) as input. KBoost uses KPCR and boosting coupled with Bayesian model averaging (BMA) to estimate the probabilities of genes regulating each other, and thereby reconstructs GRNs[10, 11, 20]. We use a greedy approach to sample the model space. That is, for every gene we build a mathematical model that predicts its expression using the kernel PCA of the expression levels of a likely subset of TFs. We then compare different KPCR TF-gene models and estimate the probability that a TF regulates a gene using BMA. The model fit is used as a proxy of the probability that a set of TFs regulate a specific gene. If there is information from other sources on likely TF targets, it can be combined with the model fit in the form of a prior. The output of KBoost is then an estimate of the probability that each TF regulates each gene.

**Fig. 2.**
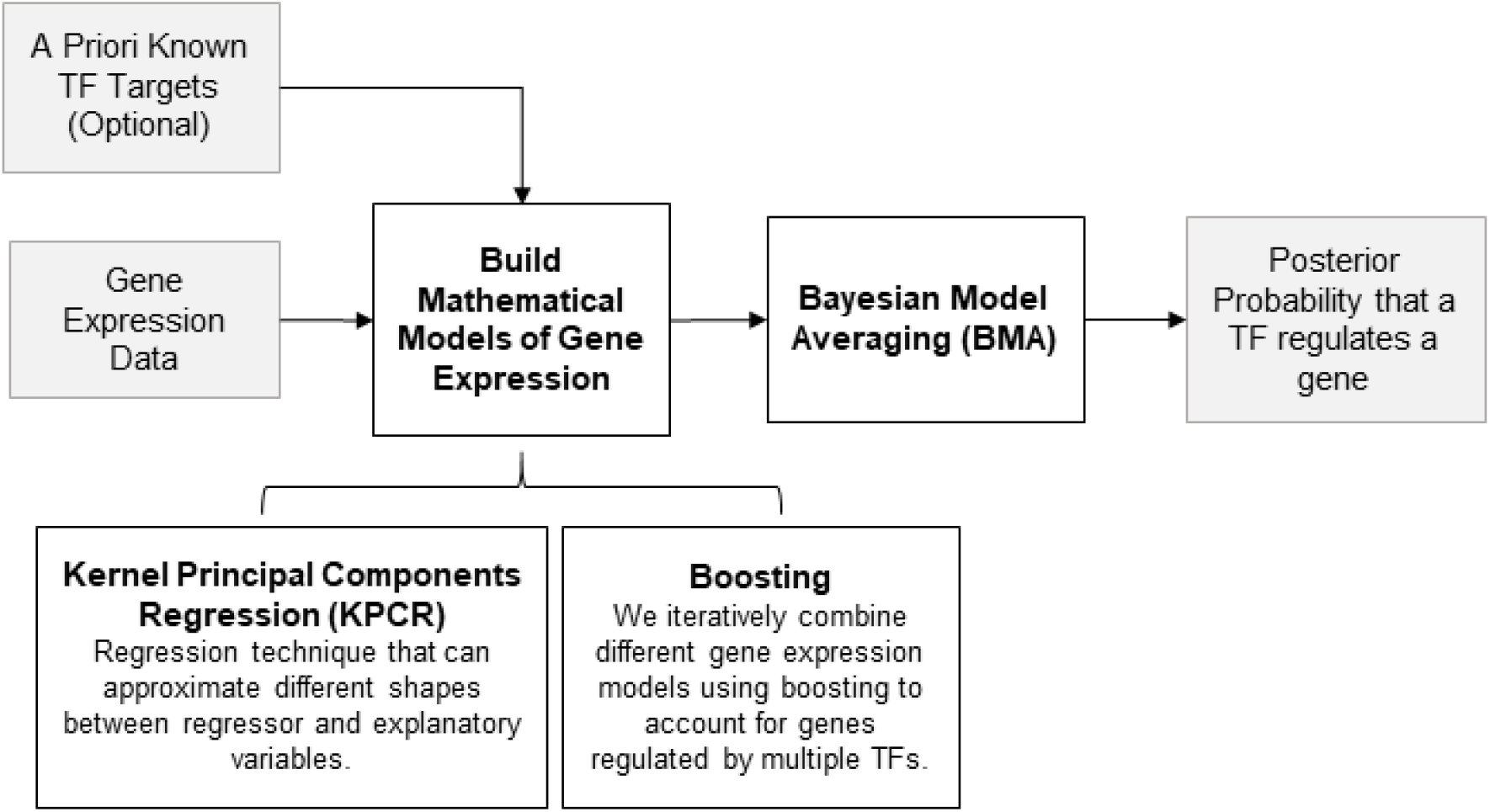
Overview of KBoost. KBoost uses KPCR in a boosting framework to infer GRNs from gene expression data. KBoost uses a list of predefined TFs and builds KPCR models to predict the expression of other genes (including TFs). These models are combined using gradient boosting. The residuals are used to estimate the probability that a TF regulates a gene.

#### 2.1.1 Modelling Gene Expression

We formulate the data as *n* independent observations ***x*** = [*x*_1_,…, *x*_*n*_]^*T*^, each with real-valued measurements of G genes: 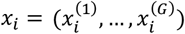. We assume that each gene’s *j* expression, 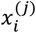, is a linear combination of non-linear functions of the expression of a subset of the gene’s TFs, noted as *A*_*j*_, plus some random noise 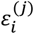, which comes from a normal distribution with zero mean and variance 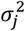:

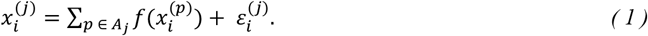

#### 2.1.1 Gene Regulatory Network Inference as Unsupervised Classification

The gene regulatory network, ***C***, would then be defined as a directed graph with *G* nodes, where each node is a gene. We write ***C*** as a *G* × *P* matrix, where *P* is the number of genes in *G* that are TFs. The entry *C*_*j,p*_ gets the value 1 if TF *p* regulates gene *j* and 0 otherwise. As many other algorithms, we formulate the problem of estimating ***C*** from ***x*** as an unsupervised classification problem, where the fit of different subsets of TFs in equation 1 is used to estimate the posterior probability that any TF regulates any gene; in other words, how well different combinations of TFs, *A*_*j*_, fit the data 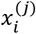. We can then consider all the combinations of TF subsets 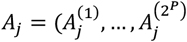, where 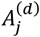 contains the integer indexes of the genes that are TFs and are part of subset *d*. The predictive probability of an element of *C*_*j,p*_ to be 1:

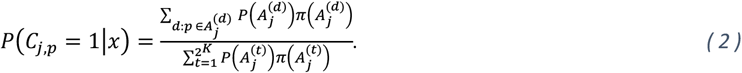

Here, 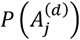 is the marginal likelihood of the model in equation 1 given the subset 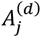, and 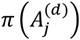 is the prior that the TFs in this subset 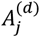 regulate gene *j*. We formulate the prior probability of a subset of TFs 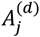 to regulate a gene, *j*, as coming from a binomial distribution:

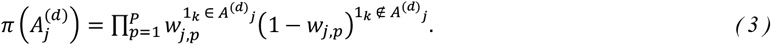

Here, the parameters of the prior are in a matrix ***W*** = [*w*_1_,…, *w*_*G*_]^*T*^ with the same dimensions as the gene regulatory network ***C***. For simplicity the marginal likelihood, 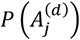, is obtained by assuming 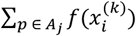 to be fixed. This assumption facilitates the inference process and reduces the computation time required in KBoost. We give the variance of the noise Jeffreys’ prior, 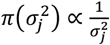. This yields the following expression for the marginal likelihood:

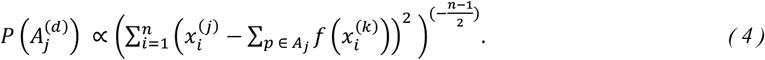

#### 2.1.3 Greedy Model Selection and Boosting

So far, we have omitted the details of two crucial tasks, first how we calculate 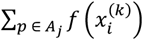, and second how we go through the different subsets 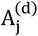. For the second task, we chose a greedy approach where we iteratively select the TF subsets with the highest posteriors 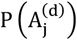 and expand them for each gene. This choice is motivated by two goals: (i) to reduce the computational burden of calculating 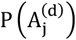 for all possible combinations of TFs, and (ii) to reduce the number of false positives. Genes that are regulated by the same set of TFs will have a high correlation between their expression levels. We assume that for each gene *j*, there are several 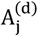 which yield a relatively high 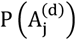 due to coregulation and thus we perform a greedy search keeping only the TFs that yield the highest 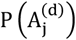 at each boosting iteration. This is similar to the LARS procedure, except that in equation (2) we include all explored models, like in the Occam Razor version of BMA [12, 21]. Therefore, equation (5) uses only the subsets *Â*_*j*_ explored by our greedy search, i.e.:

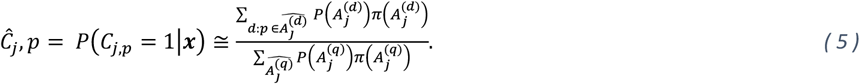

#### 2.1.4 Kernel Principal Components

Regarding the task of calculating 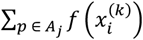, we chose to use KPCR because it has three important advantages in a boosting framework. First, KPCR is a kernel-based regression method which can approximate any potential shape that the relationship between a TF a target gene might have. We use the RBF kernel function which is the dot product of our data mapped through a nonlinear function into an infinite number of features. KPCR is a technique that allows to compute the principal components in the infinite dimensional feature space directly from the kernel matrix. A more detailed description is presented in the appendix. Second, unlike other kernel regression methods, which would need an *n* × *n* kernel matrix per TF, in KPCR we only need a few principal components per TF. This means a substantial reduction in memory usage while trying out different subsets 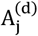. Finally, the boosting framework allows us to effectively combine principal components from kernels of different TFs. The formulation is shown below:

For each TF *p*, we first calculate the RBF kernel with width parameter *γ*:

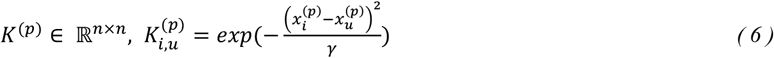

Then, we normalize the covariance matrix of the infinite dimensional features implicitly:

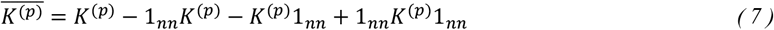

Here, 1_*nn*_ is an *n* × *n* matrix with all the values equal to 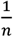. Next, we perform an eigendecomposition of 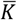:

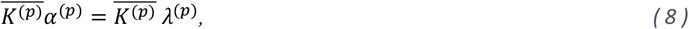

where *α* are the eigenvectors and *λ* is a diagonal matrix with the eigenvalues in descending order The first *z* KPCs are then obtained by

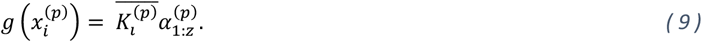

In KBoost we select KPCs by using a threshold on their corresponding eigenvalues. The KPCs are then normalized so that they have unit norm. After obtaining 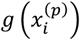, the matrices 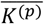 and *K*^(*p*)^ can be overwritten and thus this procedure allows us to reduce the amount of memory required. Note that the columns of 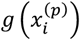 are orthogonal to each other. This means that for a model with one TF, 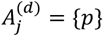, we can fit the model in equation 1 by:

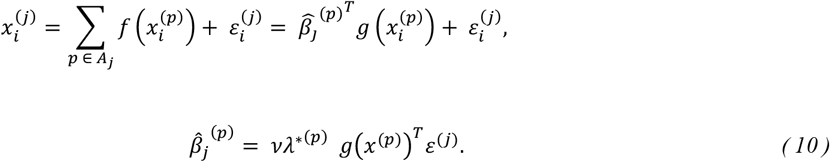

Here, 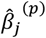 are the coefficients of the KPC regression multiplied by a constant 0 < *ν* ≤ 1, and *λ*^∗(*p*)^ is a diagonal matrix whose entries are the reciprocals of the eigenvalues that correspond to the KPCs. The constant *ν* is known as a shrinkage parameter. We propose using *ν* to counteract the effect that the number of observations *n* has on the marginal likelihood. Since the marginal likelihood changes exponentially to the power of (*n* − 1)/2, the resulting GRNs would change dramatically depending on *n*. Namely, with a large *n*, most TFs posterior probability would be close to zero while a few would be close to one. This can lead to an overestimation of the confidence we have on the predicted GRN. By shrinking the estimates of 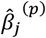, we introduce a heuristic measure that will make the resulting sum of squares less different between TF subsets 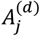, and thus reduce the effect of *n*.

#### 2.1.5 Boosting

The last equation (equation 10) shows how we fit the model in equation 1 in the case that 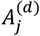 contains 1 TF. For simplicity we then follow a greedy boosting approach that is very similar to the least angle regression method. In KBoost, we first fit every single TF subset 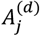 (*i*.*e*. only one TF in each model) and calculate their unnormalized posteriors. Then, we select the TF with the highest unnormalized posterior, *p*∗, and calculate its residuals, *ε*^(*j*)^. Next, we form new subsets by iteratively adding a TF, and fit the TFs to the residuals:

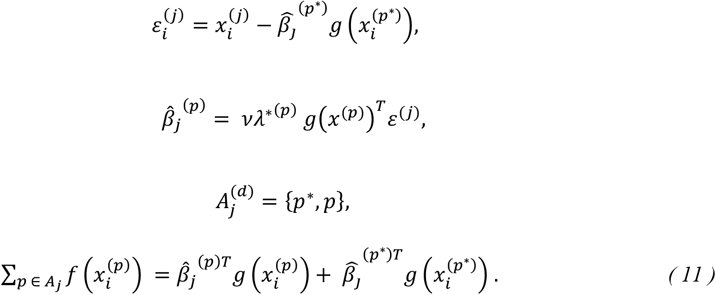

Similarly, to the previous step, we calculate their unnormalized posterior and select the TF with the highest posterior to expand next. When we use a *ν* under 1, there is a possibility that the same TF will be chosen more than once. This might reflect a TF having a very strong relationship with a particular target gene. At the end we estimate *P*(*C*_*j,p*_ = 1|*x*) by performing BMA on all explored models. In KBoost, the number of iterations represents also the maximum number of TFs per gene to be considered. The resulting network is tuned using the heuristic proposed by Slawek and Arodz [7], in which we multiply each column *p* of *P*(*C*_*j,p*_ = 1|*x*) by the sample variance of the whole column. This has the effect of preserving the sparse outdegree distribution found in most biological networks KBoost is available as an R package at: https://github.com/Luisiglm/KBoost

### 2.2 Performance Assessment

We compared KBoost against five other GRN reconstruction methods that were chosen because of their high performance in previous benchmarking exercises, such as the DREAM 4 and DREAM 5 challenges. Specifically, the algorithms included are GENIE, PLSNET, ENNET, TIGRESS, and GRNBoost2 [2, 6-8]. All algorithms were run using their default parameters, including KBoost (*if* 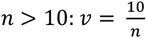, *else ν* = 0.5,, *γ* = 60 and 3 iterations). We used three different datasets to assess the comparative performance of our algorithm: IRMA [22], the *in silico* networks from the DREAM 4 multifactorial sub-challenge [2], and the DREAM 5 dataset, which contains three sets of gene expression data - one from a simulated network and two from real-life networks [3]. Before the analysis we standardized each gene’s expression to a sample variance of one and a sample mean of zero.

#### Algorithm 1. KBoost

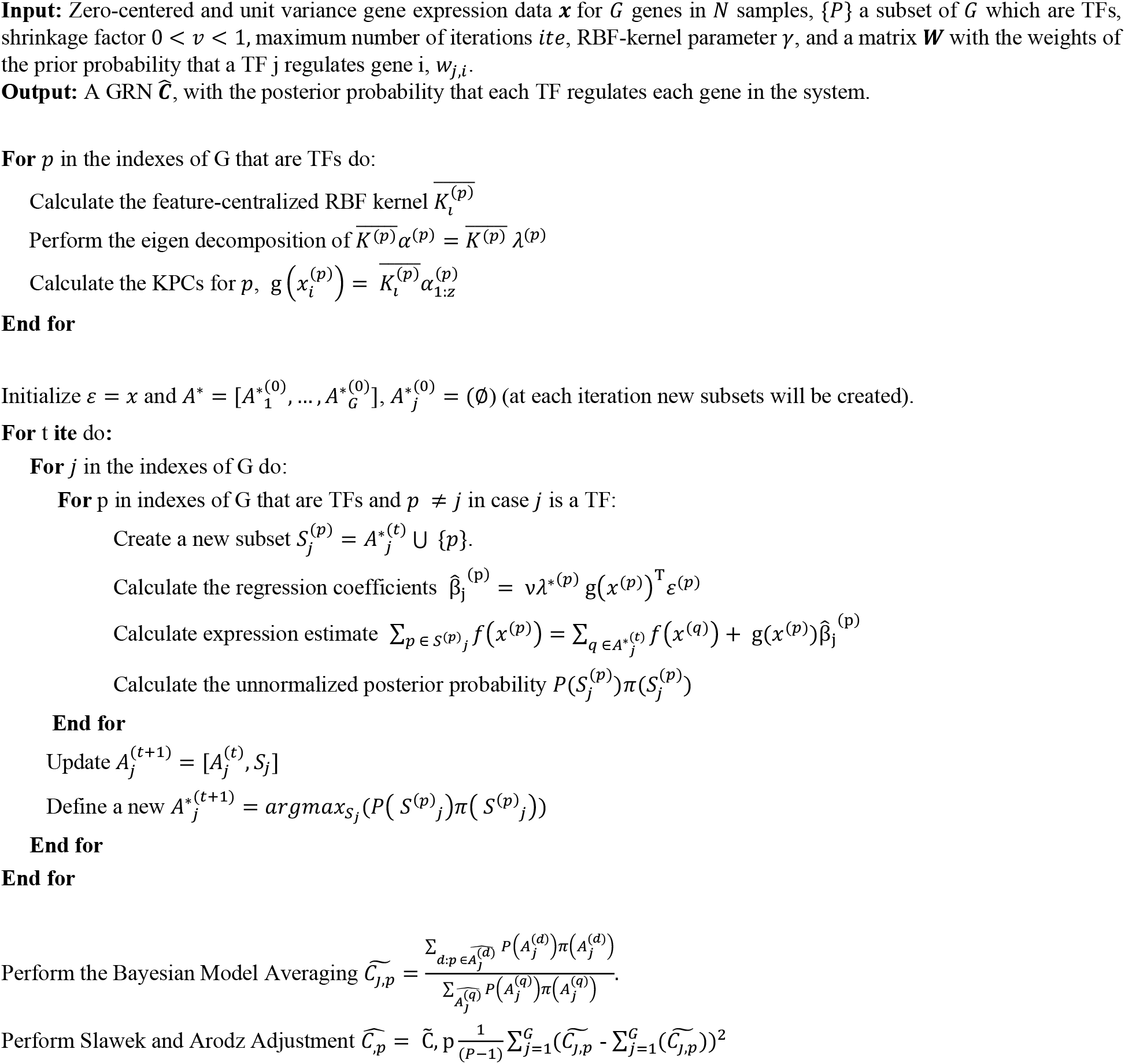

Benchmarking consisted of comparing a predicted GRN with the real GRN using common performance metrics such as the area under the precision recall curve (AUPR) and the area under the receiver operator curve (AUROC). Both methods work by changing a threshold of probability from close to 0 to 1. Regulatory interactions with a probability higher than the threshold are kept while those below the threshold are discarded. The inferred regulatory interactions are compared with the real GRN. To obtain AUPR we calculate the precision (number of correctly predicted interactions divided by the total number of predicted interactions) and the recall (number of correctly predicted interactions divided by the total number of actual interactions) at each threshold. Then, both precision and recall are plotted and the area under the curve is calculated. Similarly, for AUROC, at each threshold we calculate the recall and the false positive rate (number of incorrectly predicted interactions divided by the actual number of non-interactions). The recall is plotted against the false positive rate and the area under the curve is calculated. The AUROC value ranges from 0 to 1; 0.5 represents the number that would be achieved by random guessing and 1 is the perfect score [23]. Besides using the AUROC and AUPR, we also use the specific scores developed for the DREAM 4 and DREAM 5 challenges. The authors of the DREAM 4 and DREAM 5 challenge devised a score to benchmark different methods on their datasets. They calculated these scores using the mean of the log10 of the empirical P-value to guess a network randomly of the same accuracy as the one inferred by each specific method [3]. We used the DREAM 4 and DREAM 5 evaluation scripts to compare the different algorithms.

## 3. Results

### 3.1 Performance Assessment of KBoost

#### 3.1.1 IRMA Dataset

The IRMA dataset is a small experimental synthetic network in yeast, which is composed of five genes that regulate each other and are responsive to galactose and glucose [22]. To implement IRMA in vivo, yeast cells were transduced with expression vectors that contained five single TF promoters linked to TF encoding genes. The five genes used were ASH1, GAL80, SWI5, GAL4 and CBF1. The network can be activated by the addition of galactose and silenced by glucose. Thus, this dataset contains two small steady state sub-datasets: IRMA switch on, and IRMA switch off, respectively. The expression of the five genes was measured by q-PCR. Both IRMA switch on and IRMA switch off contain 24 observations [22]. We used the IRMA switch off dataset to select the default values of the parameters for KBoost, and these were kept fixed for the rest of the benchmark performance assessments (see supplementary data). The results show that KBoost performs similarly well in both datasets (Table 1). Furthermore, it had the highest performance in both metrics for the IRMA switch off dataset.

**Table 1.**
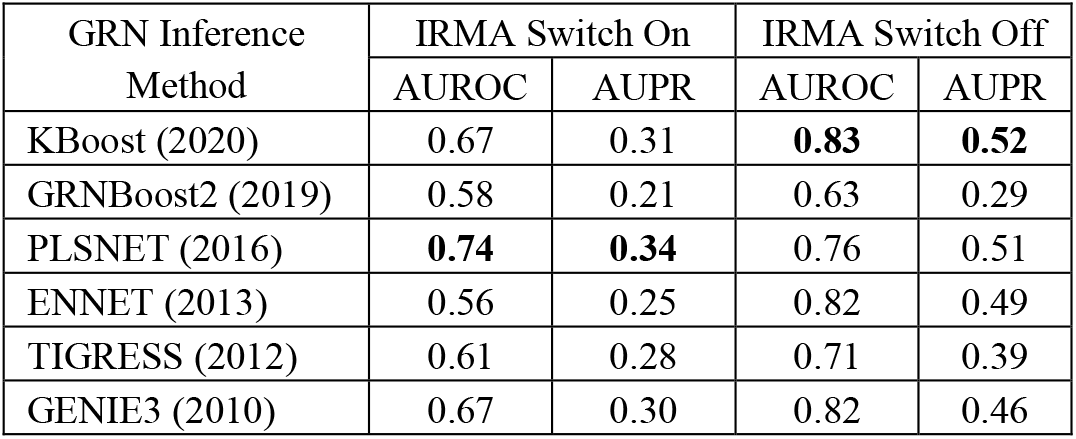
IRMA Dataset Benchmark

#### 3.1.2 DREAM 4 Multifactorial Challenge Dataset

The DREAM 4 dataset contains several different network inference challenges. We focused only on the multifactorial sub-challenge as it more closely resembles gene expression data from patients [2]. This challenge consists of 5 simulated steady-state expression profiles of 100 genes in 100 samples that were obtained by perturbing all genes simultaneously. The expression levels in the dataset correspond to steady state levels after perturbations with added normal and log-normal noise to resemble a real-life microarray experiment. The results show that KBoost, using the default parameters, generally achieved the highest performance, compared to state-of-the-art algorithms (Table 2). GENIE3, the original sub-challenge winner, had obtained a score of 37.5, whereas KBoost scores 55.93. This score is also clearly higher than that achieved by the other algorithms, which were developed after GENIE3. The results show that for several networks KBoost achieved the highest AUROC and AUPR.

**Table 2.** DREAM 4 Dataset Benchmark

| GRN Inference Method | Net1 |  | Net2 |  | Net3 |  | Net 4 |  | Net 5 |  | Score |
| --- | --- | --- | --- | --- | --- | --- | --- | --- | --- | --- | --- |
|  | AUROC | AUPR | AUROC | AUPR | AUROC | AUPR | AUROC | AUPR | AUROC | AUPR |  |
| KBoost (2020) | 0.74 | 0.15 | **0.85** | **0.31** | **0.84** | 0.25 | **0.82** | 0.27 | **0.85** | **0.32** | **55.93** |
| GRNBoost2 (2019) | 0.64 | 0.08 | 0.65 | 0.10 | 0.70 | 0.17 | 0.69 | 0.14 | 0.72 | 0.11 | 20.78 |
| PLSNET (2016) | 0.70 | 0.13 | 0.83 | 0.28 | 0.79 | 0.21 | **0.82** | 0.21 | 0.78 | 0.18 | 49.08 |
| ENNET (2013) | 0.73 | **0.17** | 0.81 | 0.26 | 0.81 | **0.29** | **0.82** | **0.29** | 0.82 | 0.28 | 52.66 |
| TIGRESS (2012) | 0.61 | 0.06 | 0.60 | 0.08 | 0.60 | 0.06 | 0.69 | 0.09 | 0.67 | 0.11 | 12.66 |
| *GENIE3 (2010) | **0.75** | 0.16 | 0.73 | 0.16 | 0.78 | 0.23 | 0.79 | 0.21 | 0.80 | 0.20 | 37.72 |

\*Winner of the DREAM 4 challenge which included 11 algorithms.

#### 3.1.2 DREAM 5 Dataset

The DREAM 5 dataset contains three sets of gene expression data, one from a simulated network (Net 1) and two from real-life networks. The three gene expression sets contain data simulated or obtained from different kinds of experiments such as drug perturbations, genetic perturbations (gene deletions or overexpression), time series sets and steady-state sets. The first network is a simulated network with 195 TFs and 1643 genes. The second network is from *Escherichia coli* with 334 TFs and 4511 genes (Net 3), and the third network is from *Saccharomyces cerevisiae* and has 333 TFs and 5950 genes (Net 4) [3]. The results show that KBoost compares favorably to most algorithms and has a similar overall performance as ENNET, a tree gradient boosting algorithm (Table 3). The results also show that most algorithms did not perform well on the experimental networks, Net 2 and Net 3.

**Table 3.**
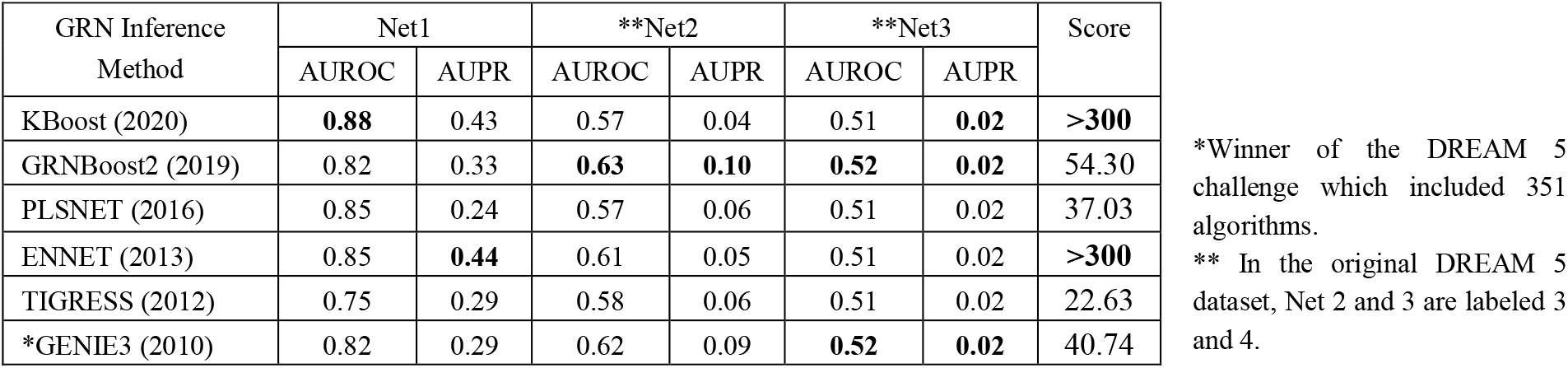
DREAM 5 Dataset Benchmark

| GRN Inference Method | Net1 |  | **Net2 |  | **Net3 |  | Score |
| --- | --- | --- | --- | --- | --- | --- | --- |
|  | AUROC | AUPR | AUROC | AUPR | AUROC | AUPR |  |
| KBoost (2020) | **0.88** | 0.43 | 0.57 | 0.04 | 0.51 | **0.02** | **>300** |
| GRNBoost2 (2019) | 0.82 | 0.33 | **0.63** | **0.10** | **0.52** | **0.02** | 54.30 |
| PLSNET (2016) | 0.85 | 0.24 | 0.57 | 0.06 | 0.51 | 0.02 | 37.03 |
| ENNET (2013) | 0.85 | **0.44** | 0.61 | 0.05 | 0.51 | 0.02 | **>300** |
| TIGRESS (2012) | 0.75 | 0.29 | 0.58 | 0.06 | 0.51 | 0.02 | 22.63 |
| *GENIE3 (2010) | 0.82 | 0.29 | 0.62 | 0.09 | **0.52** | **0.02** | 40.74 |

\*Winner of the DREAM 5 challenge which included 351 algorithms.
\*\* In the original DREAM 5 dataset, Net 2 and 3 are labeled 3 and 4.

#### 3.1.3 Running Time

The boosting framework of KBoost allows us to dramatically reduce the number of computational operations. That is because KBoost adds only one TF at a time, and the maximum likelihood solution to the regression of KPCs of only one TF does not require an inversion, since KPCs coming from one kernel are orthogonal. The resulting worst case computational complexity is governed by the eigendecomposition of the KPCA *O*(*p* · (*n*^3^)) *n* the number of observations and *p* the number of TFs. This might seem extreme but typically GRN inference datasets have at most a few hundred of observations.

We ran all algorithms on the same desktop with an Intel ® Core ™ i7-8700 CPU with 370 GHz, 6 cores and 12 logical cores, and 32 GB of RAM. The results show that KBoost is much faster for all datasets compared to all the other algorithms. The summary of all the benchmark analysis is shown in Table 4. KBoost has the best performance for all datasets except for the IRMA switch on dataset, where it performed second best. However, KBoost dramatically shortens running times, particularly for the DREAM 5 dataset.

**Table 4.** Results Summary of Running Time and Performance

| GRN Inference Method | IRMA On |  | IRMA Off |  | Total Running Time [mins] | DREAM 4 M. Challenge | Total Running Time [mins] | DREAM 5 | Total Running Time [mins] |
| --- | --- | --- | --- | --- | --- | --- | --- | --- | --- |
|  | AUROC | AUPR | AUROC | AUPR |  |  |  |  |  |
| KBoost (Matlab,2020) | 0.67 | 0.31 | **0.83** | **0.52** | **0.001** | **55.93** | **0.025** | **>300** | **8.6** |
| GRNBoost2 (Python,2019) | 0.58 | 0.21 | 0.63 | 0.29 | 0.12 | 20.78 | 0.42 | 54.30 | 68.48 |
| PLSNET (Matlab,2016) | **0.74** | **0.34** | 0.76 | 0.51 | 0.02 | 49.08 | 1.63 | 37.03 | 141.36 |
| ENNET (R, 2013) | 0.56 | 0.25 | 0.82 | 0.49 | **0.001** | 52.66 | 0.86 | **>300** | 382.46 |
| TIGRESS (Matlab,2012) | 0.61 | 0.28 | 0.71 | 0.39 | 0.06 | 12.66 | 0.65 | 22.63 | 455.77 |
| GENIE3 (Matlab,2010) | 0.67 | 0.30 | 0.82 | 0.46 | 0.003 | 37.72 | 1.60 | 40.74 | 806.75 |

#### 3.1.4 Effect of an Informative Prior Network

For several organisms, many years of research have linked certain TFs with specific genes. This information can be included into KBoost in the form of the prior ***W***. In the previous sections, we used a ***W*** with 0.5 for every of its element, giving every TF-gene relationship the same prior probability. In this section, we explore what could happen in a real-life setting, if the user had prior information from previous research studies linking certain TFs to certain genes. We used the gold standards of the DREAM 4 multifactorial dataset as prior networks and added some noise by randomly modifying different fractions of the edges. We gave the edges present in the prior networks a weight of 0.6 and the edges that were absent a weight of 0.4. The results show that even with a prior network with relatively high levels of noise (up to 50% of the edges were added or deleted from the original network) the average AUPR and AUROC are higher than when a noninformative prior was used (Fig 3). All the posterior networks had a higher AUPR than the corresponding prior networks, while for the AUROC only the prior with 5% of the edges modified had a higher AUROC in the prior than the posterior network. In general, this shows the potential to increase the accuracy and precision of the predictions by including prior information.

**Fig. 3.**
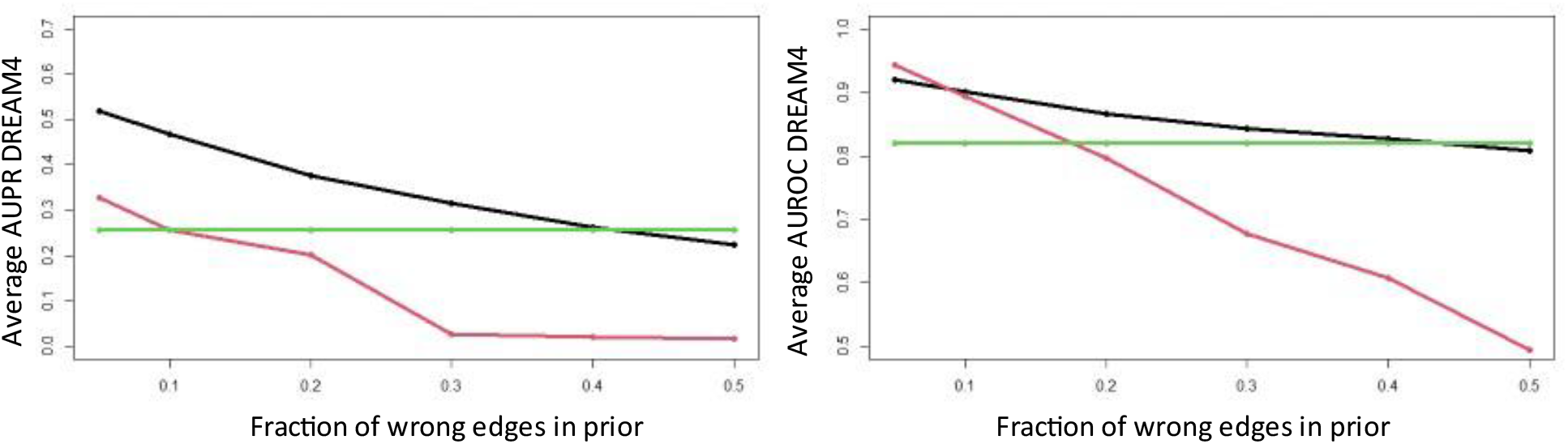
Effect of an Informative Prior into KBoost on the DREAM 4 Multifactorial Dataset. The red line represents the average of the prior networks, the green line the average results with a non-informative prior and the black line the average of the posterior networks with an informative prior. The results highlight the advantage of having prior information even if it is limited to a few TF - target gene pairs. It also shows that when the prior information is not accurate it can have a detrimental effect on performance.

#### 3.1.5 Application to the METABRIC Dataset

To examine the usefulness of KBoost to biological research we applied KBoost to the METABRIC dataset of breast cancer downloaded from cBioportal [24]. This dataset contains data for 1904 patients, of which 1898 have been stratified into subtypes. Breast cancer was one of the first cancer types where gene expression profiles were used for a molecular stratification of cancer subtypes, which correlate with disease phenotypes, prognosis and treatment response [25, 26]. Studies have identified 4 major breast cancer subtypes (luminal A, luminal B, HER2-enriched and basal), a claudin-low subtype and a normal-like group. The claudin-low is sometimes described as a particularly aggressive form of the basal subtype subtype [27] [28]. We wanted to test whether the subtypes have characteristic GRNs that can be used to distinguish them. For this, we reconstructed a network for each of the 6 subtypes using KBoost with a prior based on the GRN obtained from ChIP-Seq experiments on several TFs [29]. We gave the edges seen in the prior network a prior probability of 0.6, while for TFs in the prior network but not on our system, we gave 0.4 probability to the edges that were not observed in the ChIP-Seq. The dataset contains microarray experiments with expression levels for 24368 genes. To identify TFs we overlapped the list of 24368 genes in METABRIC with a previously published list of human TFs [30]. KBoost was ran on a desktop with an Intel ® Core ™ i7-8700 CPU with 370 GHz, 6 cores and 12 logical cores, and 32 GB of RAM. The resulting network was filtered by keeping edges with a posterior probability higher than 0.2. This threshold was found by using the F1 metric on the IRMA Off dataset. The whole dataset of 24368 genes by 1345 TFs and 1898 patients was run in 104 minutes. This is less than what most algorithms took for the DREAM 5 challenge which is significantly smaller.

The results show that KBoost reconstructed GRNs that distinguished the five main subtypes, Normal-like, Luminal A, Luminal B, HER2 enriched, and Basal/Triple negative breast cancer (TNBC) (Fig. 4 and Table 5). KBoost. We analyzed the GRNs using the closeness centrality measure, which is an indication of the number of genes a TF regulates directly or indirectly.

**Fig. 4.**
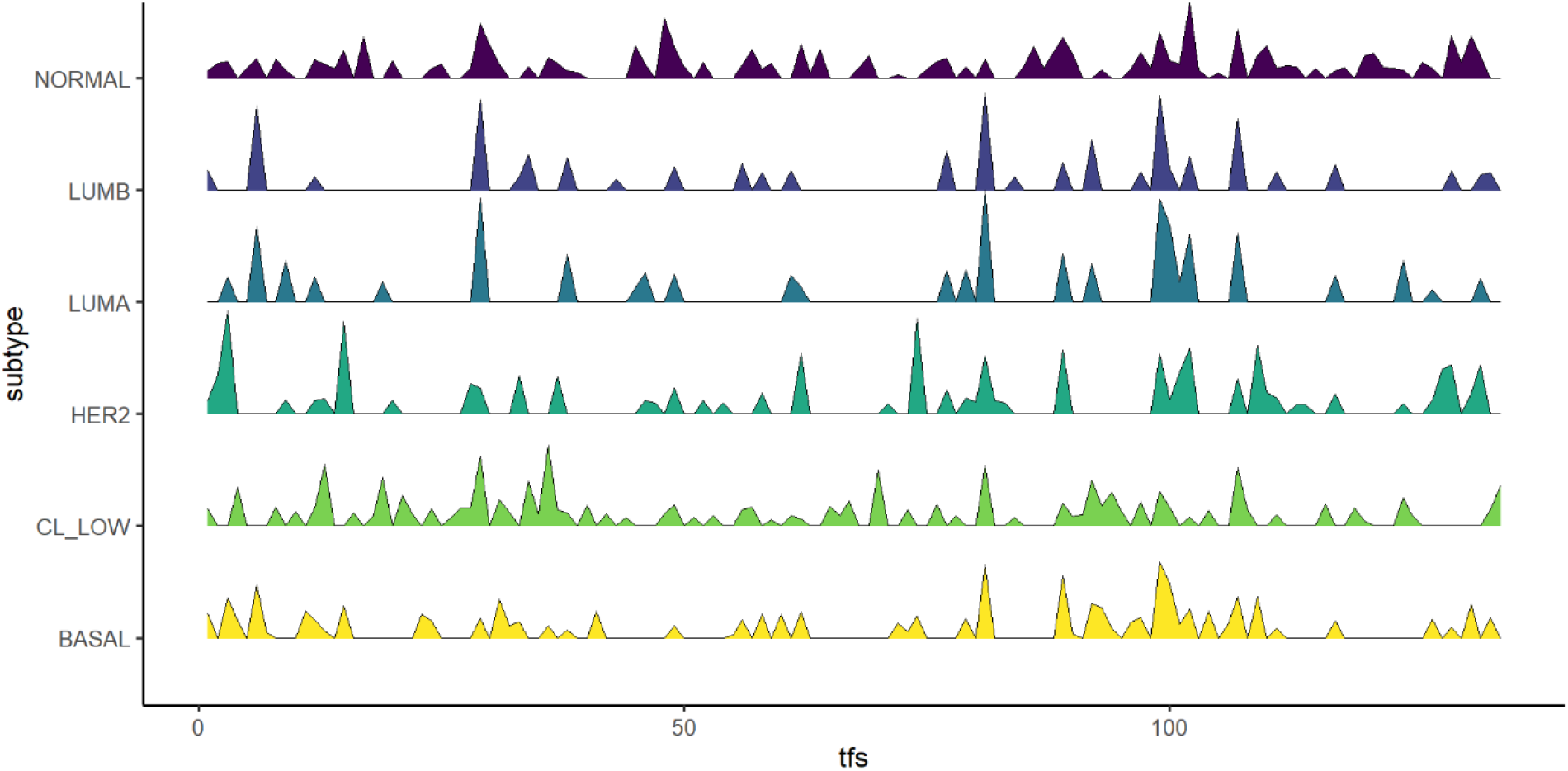
Closeness Centralities of Breast Cancer Subtypes. The peaks correspond to the closeness centrality of different TFs among the subtypes. The results show that breast cancer subtypes have differences in their underlying GRNs.

**Table 5.** METABRIC Summary

| Breast Cancer Subtype | Running Time [mins] | No. of Patients | TF Closeness Centrality Hub |
| --- | --- | --- | --- |
| Claudin Low | 13.0 | 199 | FOXM1 |
| Luminal A | 32.1 | 679 | REPIN1 |
| Luminal B | 21.6 | 461 | REPIN1 |
| HER2 enriched | 13.5 | 220 | AKAP8L |
| Basal/TNBC | 12.9 | 199 | TBX21 |
| Normal-like | 11.5 | 140 | TSC22D1 |

## Discussion

Inferring GRNs from gene expression data is an important task in bioinformatics. KBoost represents a novel algorithm that uses KPCR, boosting and BMA coupled with greedy model selection. Both boosting and BMA have been previously used in this setting. ENNET and GRNBoost2 use regression tree boosting, while TIGRESS uses a least angle regression which can be considered a form of boosting. On the other hand, BMA has mainly been used with linear regression. KPCR has advantages over using regression trees and linear regression. Both KPCR and regression trees do not need the relationship between a TF and its target gene to be linear, unlike linear regression. However, regression trees produce discontinuous functions which can be added together after several iterations to approximate a continuous function by using random forests or boosting. In contrast, KPCR directly yields continuous functions.

We show that KBoost has a very competitive performance using three different datasets. KBoost had the highest score on the multifactorial subset of DREAM 4 and the IRMA switch off dataset and tied for first place with ENNET in DREAM 5. Furthermore, though we did not use any prior information for the benchmarking, we show that in the presence of prior information the performance of KBoost can increase dramatically. In terms of running time, the advantage of using KBoost is especially clear, as it was the fastest among all algorithms across all datasets. In addition, we were able to run KBoost on the METABRIC dataset containing over 24000 genes with over 1300 TFs and over 1800 patients in less than two hours on a standard desktop computer.

We compared the GRNs from different breast cancer subtypes using the closeness centrality. Closeness centrality is a measure of a node’s importance in a network and is calculated using the number of other nodes to which it is directly and indirectly connected. Different subtypes showed differences in TF centrality. The patterns can easily be distinguished by the naked eye and correlate with increasing aggressiveness. According to the current transcriptome based classification [24, 28] the following phenotypes can be discerned: normal or normal-like is the most benign phenotype that may simply reflect normal breast tissue due to a low tumor cell content in the samples; luminal A and luminal B, which are relatively benign; HER2 enriched, which is more aggressive; and basal which has the worst prognosis. In the claudin-low subgroup, the gene FOXM1 had the highest closeness centrality. FOXM1 regulates the cell cycle and is required for the proliferation of normal breast epithelia but also promotes the proliferation of malignant cells and is often overexpressed in breast cancer [31]. The Luminal A and B GRNs were similar. In both GRNs, the TF with highest closeness centrality was REPIN1/AP4. This gene is a zinc finger protein that has been previously associated with enhanced proliferation, migration, and cisplatin resistance in breast cancer cells [32]. Interestingly, REPIN1/AP4 also was reported to reduce breast cancer cell proliferation [33], which may be related to the slow growing phenotype of luminal A/B breast cancers. The gene AKAP8L had the highest closeness centrality in HER2 enriched breast cancer. Previous studies have found AKAP8L to be associated with metastasis suppression in Luminal A, B and HER2 enriched breast cancer[34]. For the basal subtype, the gene TBX21 had the highest closeness centrality. This gene is often overexpressed in breast cancer indicating poor prognosis [37]. Finally, we found the gene TSC22D1 (TGFβ stimulated clone-22) to be important for the normal-like subtype GRN. TSC22D1 seems to have a dual role. It was identified as tumor suppressor in several cancer types including breast cancer [38]. However, it also was described that its expression correlates with tamoxifen resistance and breast cancer recurrence [39]. TSC22D1 functions in the TGFβ pathway, which can suppress tumorigenesis in early stages but promote it in later stages [40]. Together these networks highlight the gene expression differences between the breast cancer subtypes that might contribute to their observed differences in outcome. As this molecular classification of breast cancer subtypes is based on gene expression data [25, 26], it is not surprising that the subtypes differ in their GRNs. However, the identification of the TFs responsible is useful. It suggests pathogenetic mechanisms, and the dimensionality reduction associated with classifying molecular phenotypes based on GRN reconstruction facilitates the analysis and identification of potentially actionable targets.

## Supporting information

Breast Cancer Subtypes Transcription Factor Closeness Centrality

## Acknowledgements

We thank Dr. Colm Ryan for helpful discussions in the preparation of this manuscript.

