## Supplementary Data for "KBoost: a new method to infer gene regulatory networks from gene expression data"

### 1. Hyperparameter Selection for KBoost

Kboost has the hyperparameters $v,\ \gamma$ and the number of iterations for KBoost. The parameter $v$ is the shrinkage parameter for gradient boosting. It reduces the contributions of each iteration and is related to the learning rate in gradient descent. It is restricted to values between 0 and 1. The parameter $\gamma$ is the width parameter of the RBF kernel, it needs to be larger than 0. In general terms, the larger $\gamma$ the smoother the resulting regression function will be. Finally, the number of iterations controls the maximal number of TFs put together in a regression model. ChIP-Seq studies have suggested that genes have an average of 3 TFs regulating them (Gerstein, et al., 2012).

The shrinkage parameter, $v,$ in our case however it has another effect. From equation 4 and 10 the marginal likelihood is as follows:

$$x_i^{(j)} = \sum_{p \in A_j^{(d)}} f\left(x_i^{(p)}\right) + \varepsilon_i^{(j)} = \sum_{p \in A_j^{(d)}} v\hat{\beta}_j^{(p)^T} g\left(x_i^{(p)}\right),$$

$$P\left(A_j^{(d)}\right) \propto \left(\sum_{i=1}^{n} \left(x_i^{(j)} - \sum_{p \in A_j^{(d)}} f\left(x_i^{(k)}\right)\right)^2\right)^{\left(-\frac{n-1}{2}\right)}. \quad (12)$$

As mentioned in the methods section $A_j^{(d)}$ are the subet of TFs $d$ on gene $j$,$n$ the number of observations and $\sum_{i=1}^{n}\left(x_i^{(j)} - \sum_{p \in A_j} f\left(x_i^{(k)}\right)\right)^2$ is the sum of squared errors. Hypothetically speaking, if we leave the sum of squared errors divided by n fixed for two arbitrary subsets $A_j^{(d)}$ and $A_j^{(d+1)}$, where the sum of squred errors is larger in subset $d+1$ than $d$, the ratio $P\left(A_j^{(d)}\right) / \left(P\left(A_j^{(d+1)}\right) + P\left(A_j^{(d)}\right)\right)$ becomes closer to 1 as $n$ increases while $P\left(A_j^{(d+1)}\right) / \left(P\left(A_j^{(d+1)}\right) + P\left(A_j^{(d)}\right)\right)$ becomes closer to zero. This is easier to see in log form:

$$\log\left(P\left(A_j^{(d)}\right)\right) = Q + \left(-\frac{n-1}{2}\right)\log(sse_d) + \left(\frac{n-1}{2}\right)\log(n)$$

$$sse_d = \sum_{i=1}^{n} \frac{\left(x_i^{(j)} - \sum_{p \in A_j^{(d)}} f\left(x_i^{(k)}\right)\right)^2}{n}$$

Here, Q is a constant, let $sse_d$ and $sse_{d+1}$ be the sum of squared errors divided by n for $A_j^{(d)}$ and $A_j^{(d+1)}$ respectively. The ratio increases:

$\log\left(P\left(A_j^{(d)}\right)\right) - \log\left(P\left(A_j^{(d+1)}\right)\right) = -\left(\frac{n-1}{2}\right)\log(sse_d) + \frac{(n-1)}{2}\log(sse_{d+1}) = \left(\frac{n-1}{2}\right)\left(\log(sse_{d+1}) - \log(sse_d)\right).$

This means that for two models with the same fit, depending on n we can observe very different Bayesian model averages. If n is sufficiently large and $sse_d$ is lower than $sse_{d+1}$, then $P\left(A_j^{(d+1)}\right) \ll P\left(A_j^{(d)}\right)$ yielding the result $P\left(A_j^{(d+1)}\right) + P\left(A_j^{(d)}\right) \cong P\left(A_j^{(d)}\right)$. The Bayesian model averages under this conditions would result in 1 for $A_j^{(d)}$ and 0 for $A_j^{(d+1)}$. In our case, we are using BMA as an estimate to the probability that a TF regualtes a gene. In conditions as those described above using a shrinkage parameter, we could overestimate the confidence of our predictions. As a heuristic, we propose reducing $\nu$ in the boosting algorithm to balance this. If $\nu$ decreases as a funnction of $n$, then the difference $\log(sse_{d+1}) - \log(sse_d)$ would be less dramatic because the contribution of the TF models are reduced. This can be seen in supplementary figure 1. Furthermore, since we use the same shrinkage parameter for all models, the model ranking remains identical when using 1 boosting iteration.

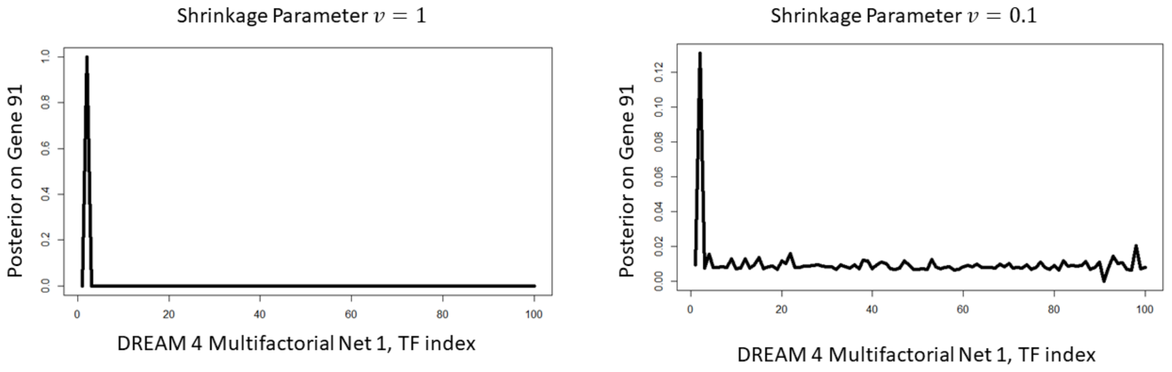


**Figure S1. Effect of the Shrinkage Parameter on the Posterior.** We ran KBoost on the DREAM4 multifactorial network 1, using $v$ of 1 and 0.1 with 1 iteration and a parameter $\gamma$ of 60. The results highlight the effect of the shrinkage parameter at reducing the disparity between models which is a product of a large $n$.

To see how different parameters affected the performance of KBoost, we ran KBoost on the IRMA off dataset with different combinations of values for each parameter and used the AUPR and AUROC metric to asses its performance. We fixed the number of iterations to 3, as they represent the maximum number of TF to be considered to regulate a gene together. For the other parameter we used $v$ values ranging from 0.0001 to 1 and $\gamma$ values from 1 to 100. We focused on finding ranges of values in which both metrics were consistently high rather than a single value that had the maximum as it could be an artifact specific to this dataset. The results are shown in supplementary figure 2. The performance seemed to plateau for large widths $\gamma > 50$ given shrinkage parameters $v$ under 0.5.

Interestingly, the behavior is similar in the five networks of the DREAM 4 dataset (Figure S3). While the shrinkage parameter, $v$, has an effect on the sparsity of the posterior, and the number of iterations corresponds to the maximum number of TFs per, the width parameter, $\gamma$, can be associated with overfitting. An RBF kernel regresion model with a low width $\gamma$ will be able to fit every point of the data exactly, this increases the chance of fitting spurius patterns. We investigated the effect of $\gamma$ in the network 1 of the DREAM 4 multifactorial challenge dataset. We performed a three fold cross validation with a grid of iteration from 1, 3, 5, 10 and 20, shrinakges from 0.001, 01, 0.3 0.5 and 1 and widths, $\gamma$, taking the values of 0.1, 10 , 20 , 60 and 100 . The results indicate in all cases that $\gamma \geq 10$ had a lower sum of squared errors in the validations set (Figure S4).

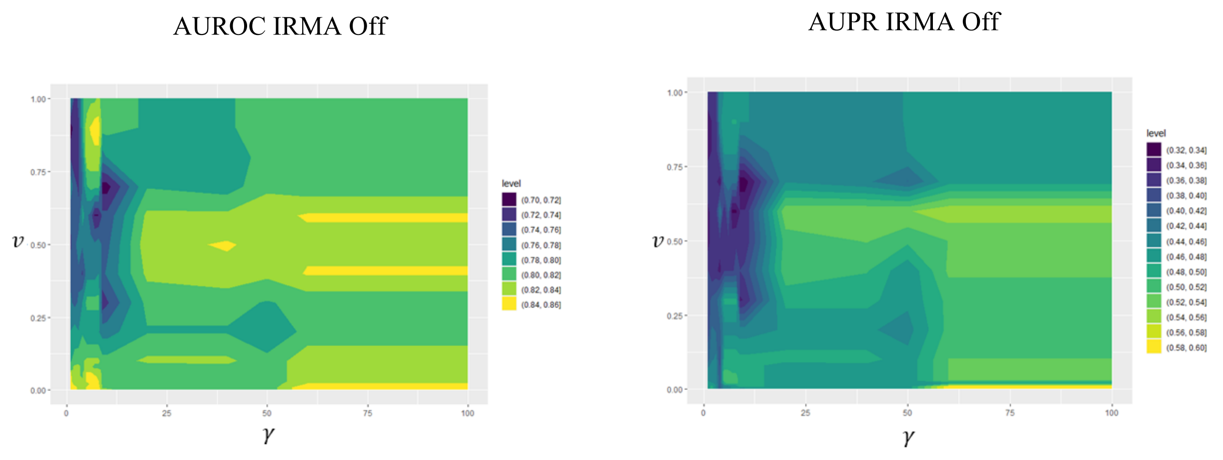


Figure S2. Hyperparameter Selection for KBoost

The results show the effect of the width and shrinkage parameters on the performance on the IRMA Off dataset. The performance seems to plataeu for large widths $\gamma$ for any value of $v$ . We fixed the number of iterations at 3, assuming a maximum of 3 TFs per gene.

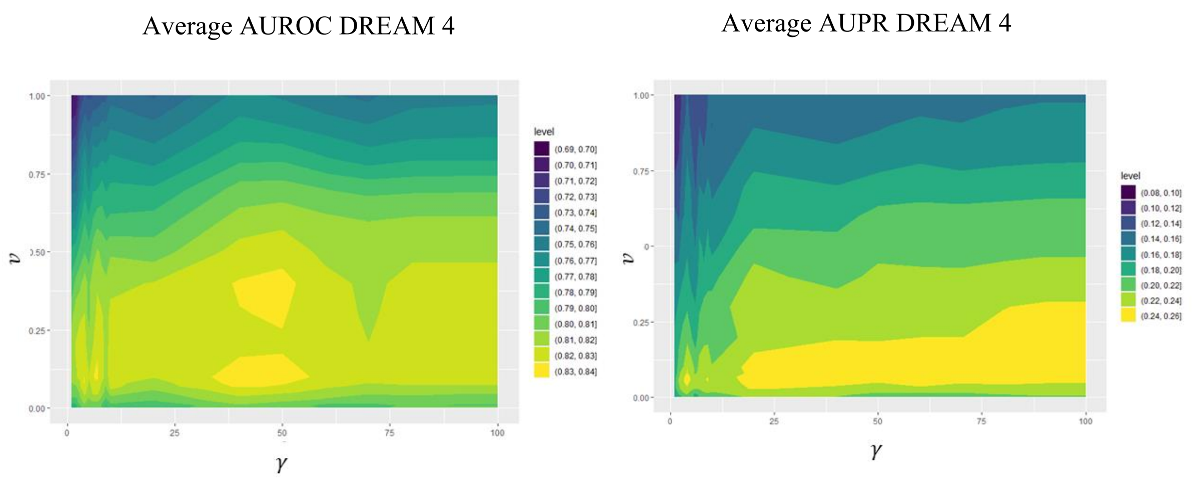


Figure S3. Hyperparameter Effect for KBoost on the DREAM 4 Multifactorial Dataset.

The results show the effect of the width and shrinkage parameters on the performance on the DREAM4 multifactorial dataset. The performance seems to plataeu for large widths $\gamma$ for relatively low values of $v$. We kept the number of iterations fixed at three.

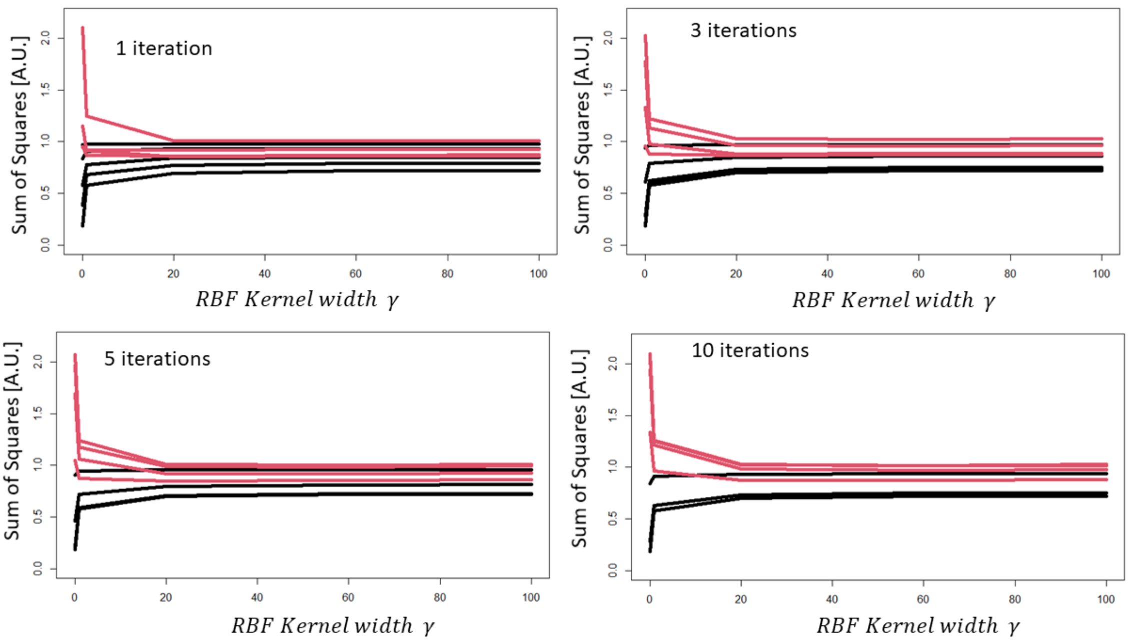


Figure S4. RBF Kernel Width Cross-Validation on Network 1 of DREAM 4

In all cases, the red line corresponds to the average sum of squared errors in the validation set. Each line in either colour corresponds to the results obtained at a shrinkage parameter, $v$, of 0.01, 0.1, 0.3, 0.5 and 1.0. The results show that at the width parameter $\gamma$ values that show the highest performance there is less overfitting and a relatively lower sum of squared errors on the validation set.
